# NGseqBasic - a single-command UNIX tool for ATAC-seq, DNaseI-seq, Cut-and-Run, and ChIP-seq data mapping, high-resolution visualisation, and quality control

**DOI:** 10.1101/393413

**Authors:** Jelena Telenius, The WIGWAM Consortium, Jim R. Hughes

## Abstract

With decreasing cost of next-generation sequencing (NGS), we are observing a rapid rise in the volume of ‘big data’ in academic research, healthcare and drug discovery sectors. The present bottleneck for extracting value from these ‘big data’ sets is data processing and analysis. Considering this, there is still a lack of reliable, automated and easy to use tools that will allow experimentalists to assess the quality of the sequenced libraries and explore the data first hand, without the need of investing a lot of time of computational core analysts in the early stages of analysis.

NGseqBasic is an easy-to-use single-command analysis tool for chromatin accessibility (ATAC, DNaseI) and ChIP sequencing data, providing support to also new techniques such as low cell number sequencing and Cut-and-Run. It takes in fastq, fastq.gz or bam files, conducts all quality control, trimming and mapping steps, along with quality control and data processing statistics, and combines all this to a single-click loadable UCSC data hub, with integral statistics html page providing detailed reports from the analysis tools and quality control metrics. The tool is easy to set up, and no installation is needed. A wide variety of parameters are provided to fine-tune the analysis, with optional setting to generate DNase footprint or high resolution ChIP-seq tracks. A tester script is provided to help in the setup, along with a test data set and downloadable example user cases.

NGseqBasic has been used in the routine analysis of next generation sequencing (NGS) data in high-impact publications ^1,2^. The code is actively developed, and accompanied with Git version control and Github code repository. Here we demonstrate NGseqBasic analysis and features using DNaseI-seq data from GSM689849, and CTCF-ChIP-seq data from GSM2579421, as well as a Cut-and-Run CTCF data set GSM2433142, and provide the one-click loadable UCSC data hubs generated by the tool, allowing for the ready exploration of the run results and quality control files generated by the tool.

**Availability:** Download, setup and help instructions are available on the NGseqBasic web site http://userweb.molbiol.ox.ac.uk/public/telenius/NGseqBasicManual/external/

Bioconda users can load the tool as library “ngseqbasic”. The source code with Git version control is available in https://github.com/Hughes-Genome-Group/NGseqBasic/releases.

## INTRODUCTION AND RESULTS

Advances in next generation sequencing (NGS) techniques allows for the production of vast amounts of genomics sequence data sets. To manage and prioritise these data sets, it is vital to provide a fast and reliable checkpoint after the performance of the experiments, to determine the success and quality of the data, before either investing in in-depth analysis of the data or returning to optimise experimental conditions. However, such analysis would traditionally require extensive bioinformatics experience and so be performed by a dedicated computational scientist, considerably slowing down this process. To speed up the feedback between experimental data generation and analysis an approach is needed capable of performing extensive automated, read mapping and filtering, and should generate the appropriate quality control reports and data metrics, but require only standard computational experience.

In this way the analysis results should be able to guide the experimentalist in improving the experimental protocol (if data quality was poor) or provide sufficient confidence to allow for further data generation, freeing the time of the bioinformatician for more downstream in-depth analysis, as well as guide in the potential troubleshooting of the sample mapping and trimming. This “checkpoint data” should be easy to share between group members and collaborators, and be presented in an easy-to-browse format, including visualisations (readily comparable to publicly available data sets), and accompanied with quality metrics and tool log files to provide transparent analysis. Preferentially, this analysis should be provided as a single-command tool, easy enough for a basic level UNIX user to run, but with a comprehensive panel of run options to adjust run parameters for fine tuning by more experienced users.

Many of the currently available chromatin accessibility and ChIP sequencing analysis pipelines generate the output data to make such a “checkpoint data set” ^3–8^. However, they do not collect the data to a single-click loadable visualisation, including all the metrics and analysis reports (for tool comparisons, see Supplementary Table 1) and therefore are more time consuming and require higher levels of computational expertise of the user. Other solutions provide GUI-based analysis, where modifying the source code is more time consuming^9–11^. Promising database-based all-inclusive data analysis platforms have been developed ^12,13^, but these are feasible for only large core facilities, where the time and effort necessary for system administration and database management becomes cost-effective.

To address this lack of a generally applicable platform we have built NGseqBasic, a single-command pipeline that is easy to setup and with a comprehensive and easy to interpret output. NGseqBasic includes all standard analysis and QC steps and supports the most commonly used parameters of the best practises of mapping and trimming tools and has a rich pallet of option that allow for bespoke analysis.

NGseqBasic produces within a single run the mapping, trimming, short read rescue, quality control, duplicate filter, peak call, footprint, and visualisation of the data (Fig. 1), and generates a one-click loadable UCSC data hub ^14^ (Fig 2.) Integrated into this hub is a linked html description page of the mapping statistics and run log files for all steps (Fig 3.). This data hub can be readily viewed with existing publicly available UCSC data, and shared with collaborators simply by distributing the data hub web address. Various parameters are provided for the user to fine-tune the analysis (for list of available parameters, see Supplementary table 2). All the output data files are stored in the run folder, readily available for further downstream analysis of the data set. NGseqBasic has been used in several peer-reviewed publications^1,2,15–19^, is actively developed, and accompanied with a version control repository. It has a wide user base, reaching ~1000 pipeline runs per annum, ~20% of which is independent installations outside the institute it was developed in.

**Figure 1.**
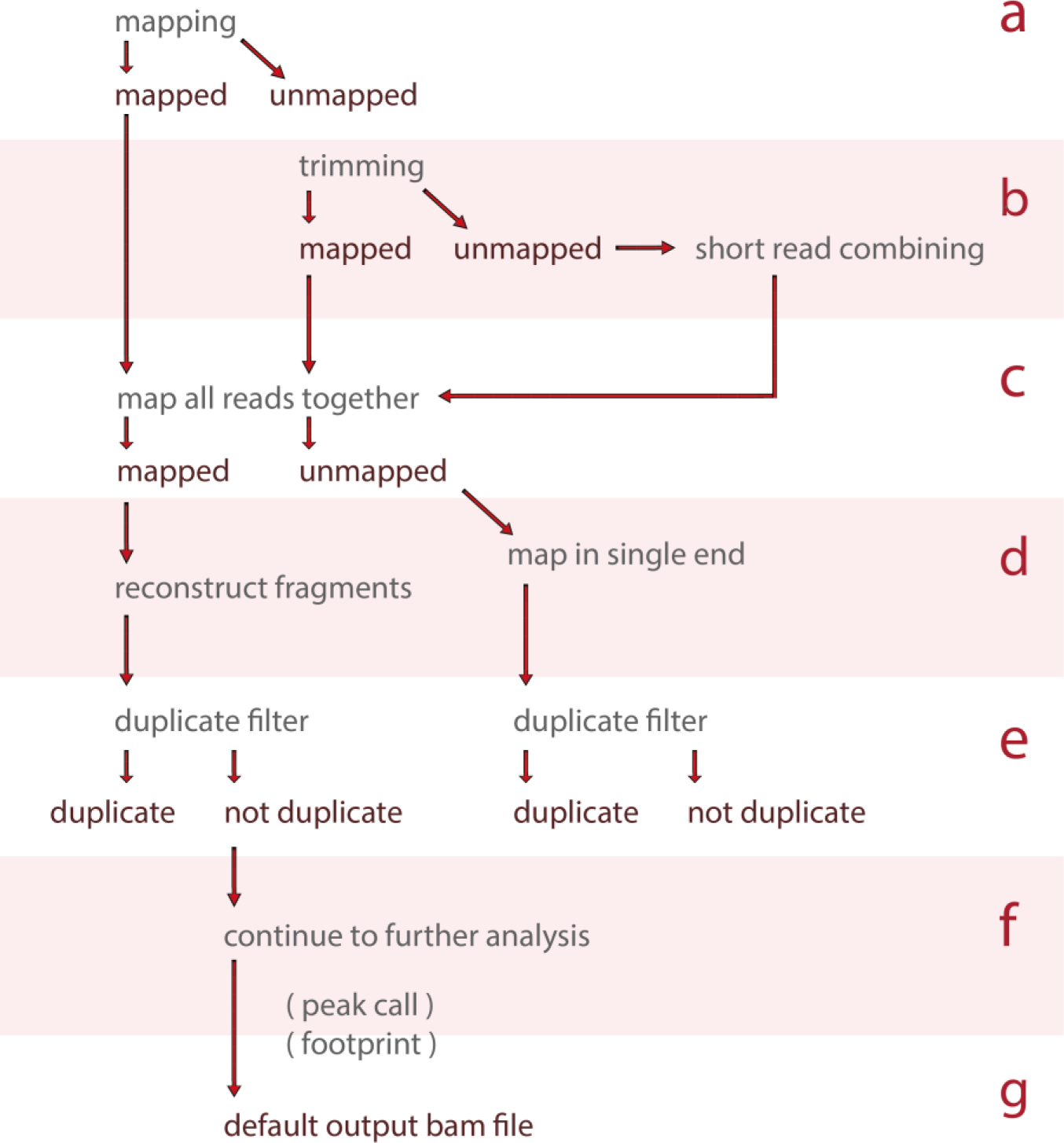
NGseqBasic data flow. Data flow steps a) - e) are accompanied with run log files and quality control reports. Trimming and short read combining in step b) can be skipped. Visualisations are generated from all data in step e). For genomes which have blacklist regions provided (mm9, mm10, hg18, hg19), filtering contains also removal of blacklisted regions. Peak call and footprint are generated in step f). Default output is given in step g), and all data from step e) can be requested as additional bam output files.

**Figure 2.**
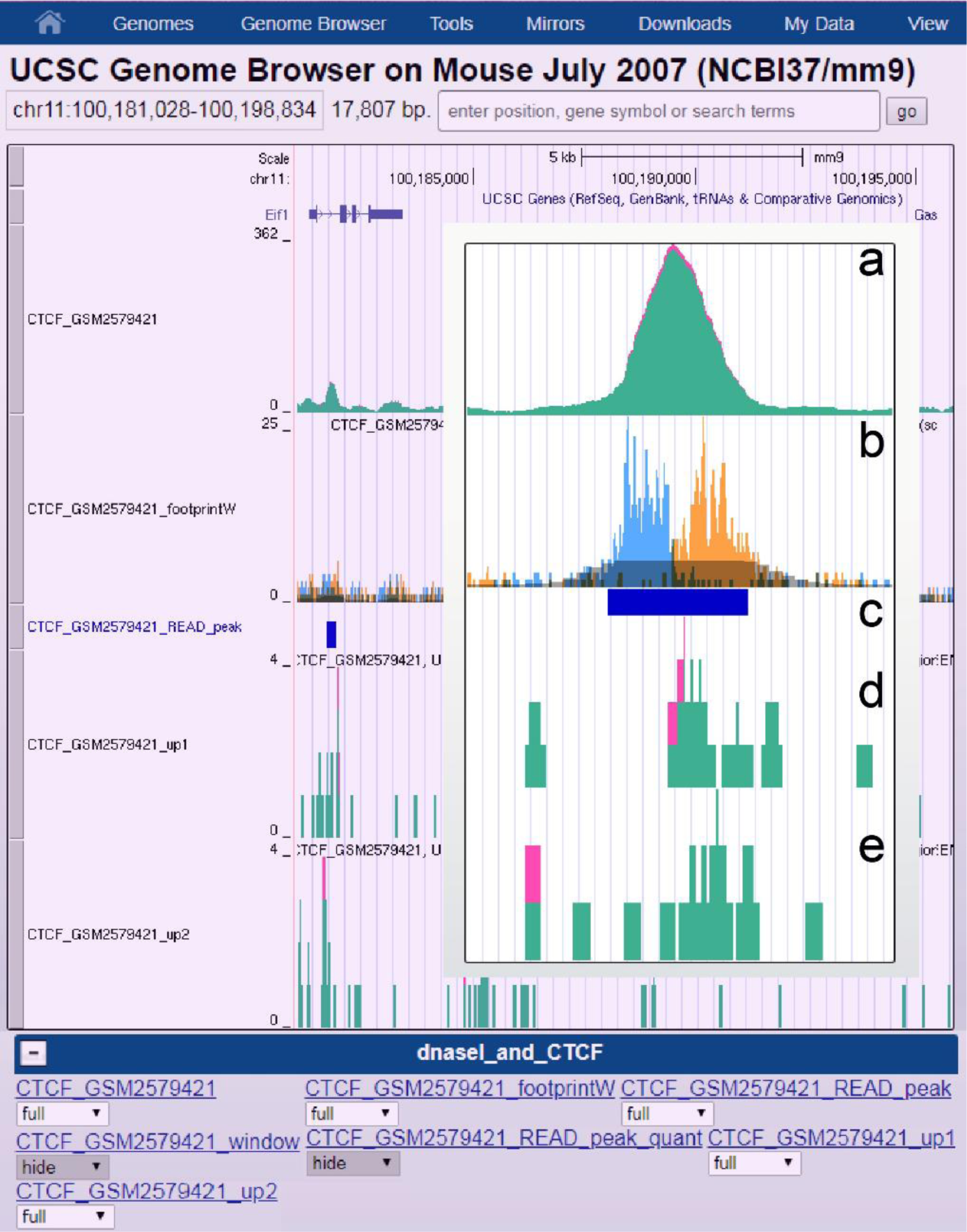
NGseqBasic produces single-click loadable UCSC data hub. The data hub address can be copypasted into UCSC data hub load field to load the hub. The tracks will be readily visible. From top down: a) All mapped reads (red), duplicate- and blacklist-filtered reads (green), b) Footprints of the data (leftmost 1 base of each read pair in cyan right-most 1 base in orange. Grey shade is down-scaled filtered mapped reads from a). Peak call of the data is shown in c), and only-single-end mapping reads are plotted in the bottom of the figure: read1 in d) and read2 in e).

**Figure 3.**
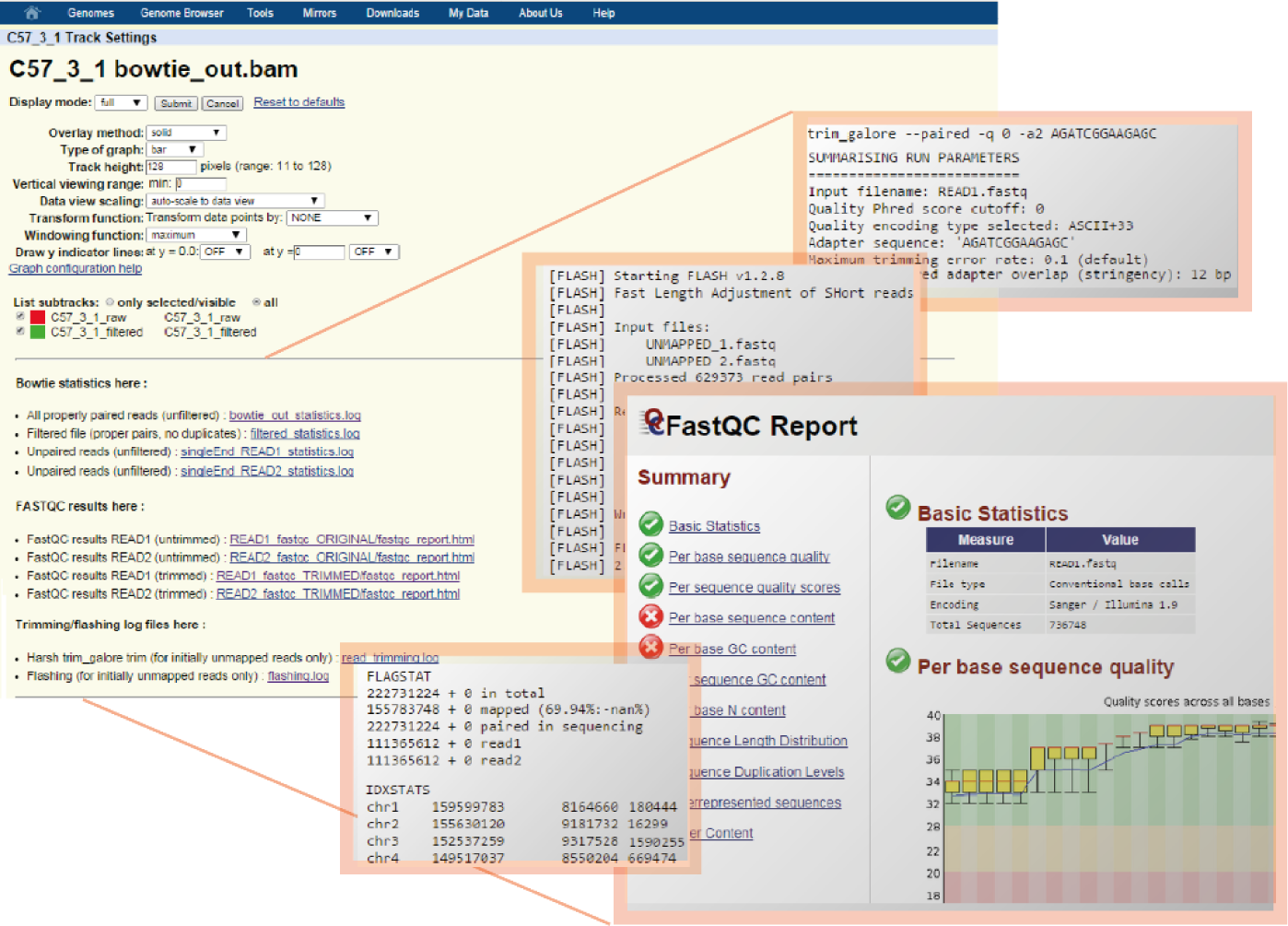
NGseqBasic stores the run log files and quality control reports in a html page accessible via the UCSC data hub. Clicking or right-clicking any of the tracks in the UCSC browser, will open the “track settings” page, where all the reports are stored as html links. From top down: trimming report (trim_galore output log), short read combining report (flash output log), FastQC report, samtools report (idxstat, flagstats). All analysis stages and output files are documented in the output logs.

Additionally, NGseqBasic has integrated analysis modes to increase the resolution at which the data can be visualised and interpreted. DNase-seq data has been shown, if analysed correctly, to contain high resolution “footprints” of transcription factor binding within the open chromatin regions^20^. Similarly, it has been shown that the independent visualisation of the left and right hand reads in a paired end ChIP-seq experiment more precisely localises the bound protein^21^. To provide these high-resolution data NGseqBasic plots the distribution of fragment ends which it derives from the aligned sequences. This Footprint visualisation is included in the analysis hubs and reveals the regions of DNA protected from DNase digestion by bound protein factors, and the high-resolution binding site of the protein, for DNase-seq and ChIP-seq data, respectively.

We used NGseqBasic to analyse DNaseI-seq data from GSM689849, and CTCF-ChIP-seq data from GSM2579421 (Figure 4) using these high resolution analysis modes for DNase-seq and ChIP-seq. NGseqBasic was able to plot the data in a clear and illustrative manner, such that the DNaseI and CTCF footprint tracks readily determine in near basepair-resolution the exact location of the CTCF binding sites.

**Figure 4.**
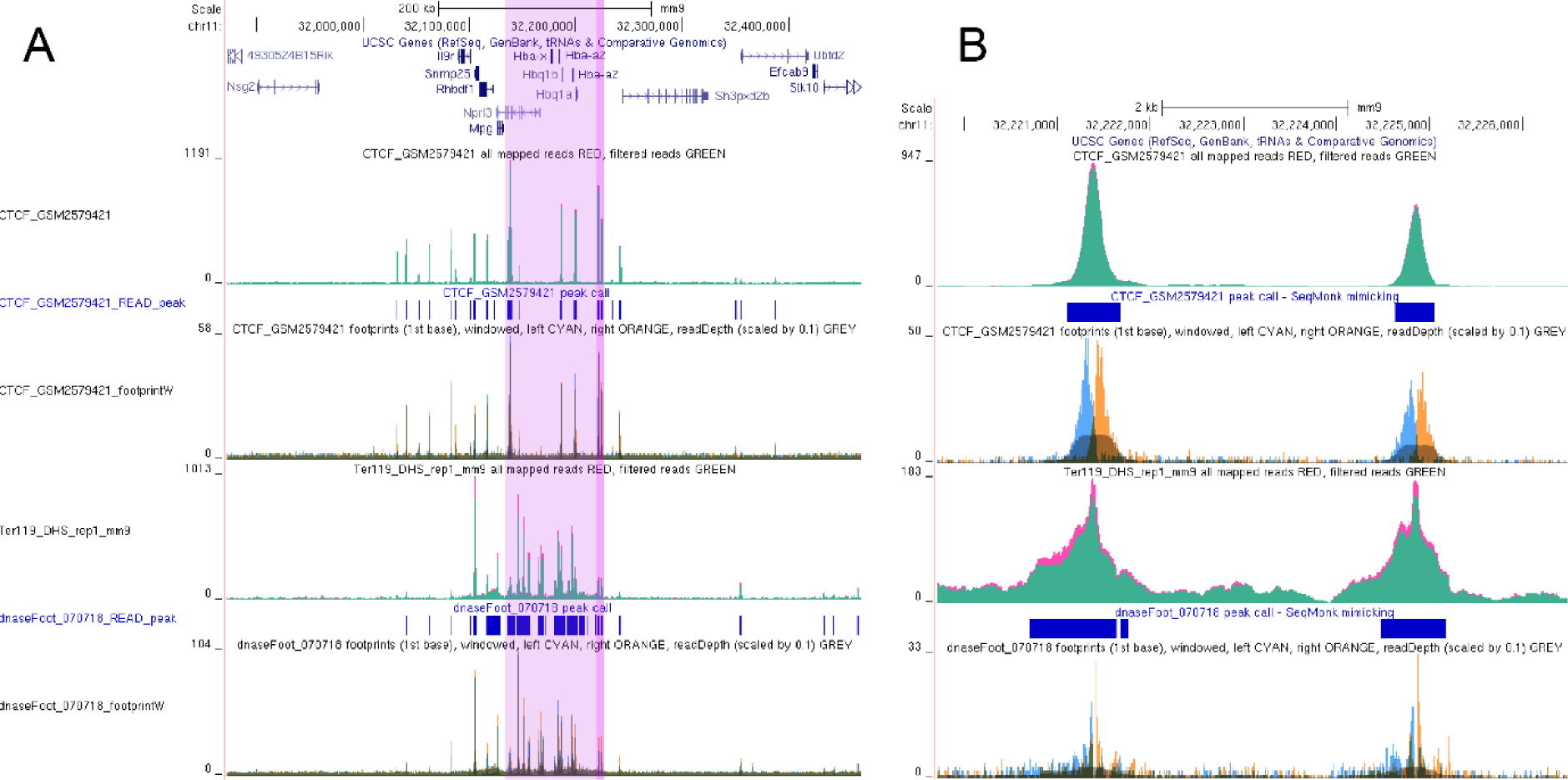
NGseqBasic data hub screenshot for mouse erythroid cell samples GSM689849 (DNaseI) and GSM2579421 (CTCF). A) Alpha globin locus control region in light purple, and its right-most CTCF cites in purple highlight. The tracks (from the top): mapped CTCF read pairs, CTCF peak call (default parameters), CTCF footprints (left- and rightmost 1base of the mapped read pairs), mapped DNaseI read pairs, DNaseI peak call (default parameters), and DNaseI footprint. B) The same data, zoomed into the right-most CTCF cites of the locus control region (highlighted in purple in A), to show (from the top): mapped CTCF read pairs, CTCF peak call (default parameters), CTCF footprints (left- and rightmost 1base of the mapped read pairs), mapped DNaseI read pairs, DNaseI peak call (default parameters), and DNaseI footprint. The data hub addresses and run parameters are given in Table 1.

Similar to DNAse-seq the Cut-and-Run ChIP-seq method, which employs the MNase nuclease, is also capable of producing footprints for DNA bound proteins. To this end we used NGseqBasic to analyse the published CTCF dataset (GSM2433142) and reproduced the original Cut-and-Run publication^22^ figure 6A lower panel results for (Figure 5). The NGseqBasic -generated data hub can readily illustrate the observation made by the original Cut-and-Run authors^22^, that by restricting inspection to fragment sizes 120 bases or smaller (fragments within one nucleosomal distance), the actual CTCF binding site is better resolved (Figure 5A). The exact location of the CTCF binding site can be determined with NGseqBasic footprints (the left- and right-most 1 base of the read pairs), when inspecting the fragments within one nucleosomal distance (fragment size 120 bases or smaller) (Figure 5B).

**Figure 5.**
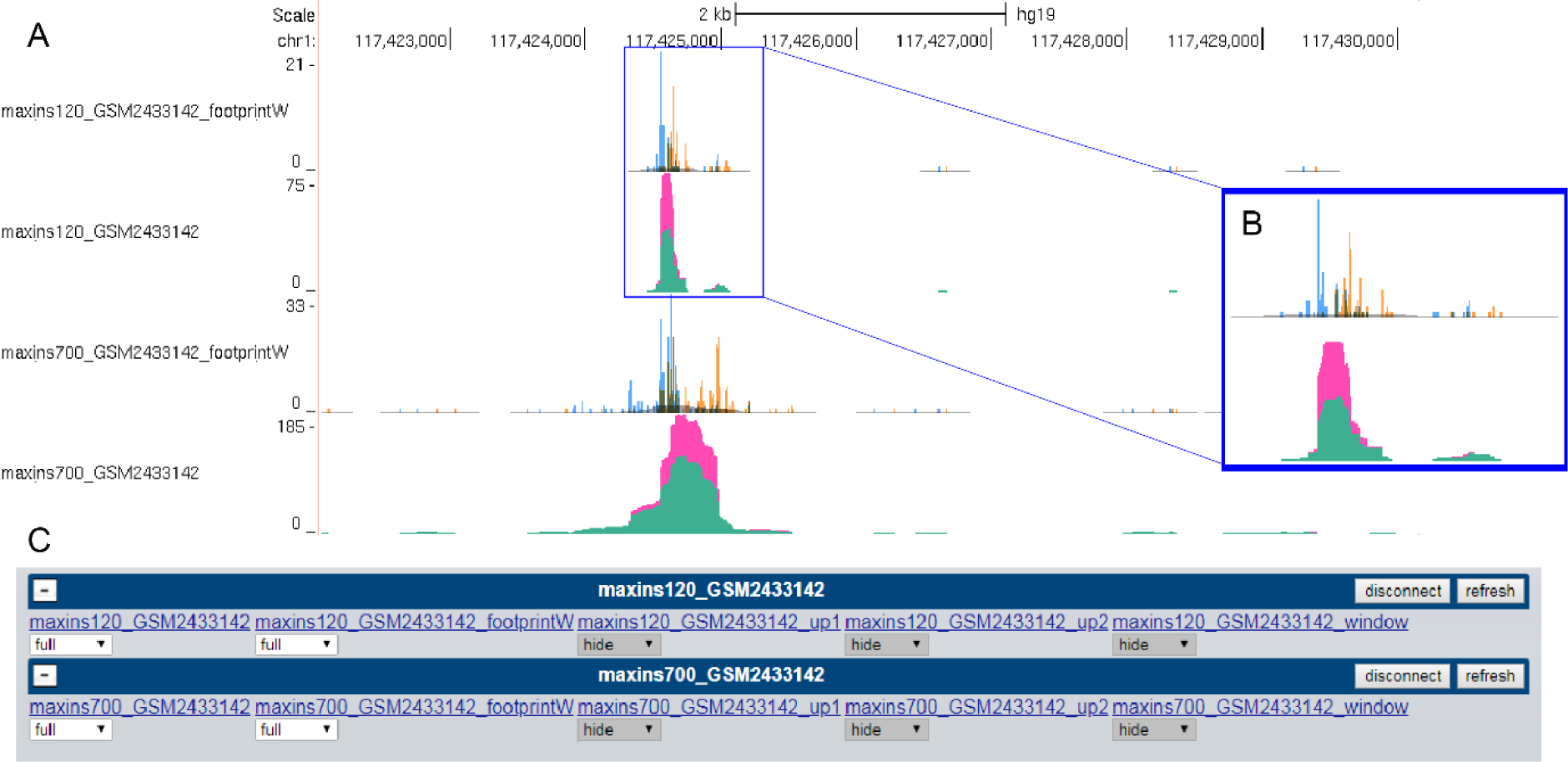
Screenshots from the UCSC data hub, generated by NGseqBasic, when CTCF Cut-and-Run data set GSM2433142 was ran, to reproduce the original Cut-and-Run publication^22^ figure 6A lower panel results. Two bowtie2 insert size parameters were used in the run: --maxins 120, and --maxins 700. The data hub addresses and run parameters are given in Table 1. 5A) NGseqBasic -generated data hub can readily illustrate the observation made by the original Cut-and-Run authors^22^, in figure 6A lower panel, that by restricting inspection to fragment sizes 120 bases or smaller (fragments within one nucleosomal distance), the actual CTCF binding site is better resolved. The footprints (left- and right-most 1bp of each read pair) are shown in the NGseqBasic data hub as cyan and orange tracks, respectively. 5B) The inset further illustrates, how the CTCF binding site, residing between the left and right peaks can be determined with high accuracy in NGseqBasic results, when inspecting the fragments within one nucleosomal distance (fragment size 120 bases or smaller). 5C) The UCSC data hub generated by NGseqBasic allows the user to flexibly choose the tracks visualised in the browser window, by grouping the tracks into a “data hub” unit, which is visible in the main browser window, alongside the UCSC provided public annotation tracks.

**Table 1.** List of UCSC data hubs generated for the test data sets.

|  |
| --- |
| DNaseI (GSM689849) + CTCF (GSM2579421) data hub |
| http://userweb.molbiol.ox.ac.uk/public/telenius/NGseqBasicManual/external/GEODataHubs/dnaseI_and_CTCF/HubFolder/hub.txt |
| CTCF Cut-and-Run (GSM2433142) data hubs |
| Bowtie2 --maxins 120 |
| http://userweb.molbiol.ox.ac.uk/public/telenius/NGseqBasicManual/external/GEODataHubs/maxins120_GSM2433142/HubFolder/hub.txt |
| Bowtie2 --maxins 120 |
| http://userweb.molbiol.ox.ac.uk/public/telenius/NGseqBasicManual/external/GEODataHubs/maxins700_GSM2433142/HubFolder/hub.txt |
| Instructions to load the data hub to UCSC server : |
| NGseqBasic hub load tutorial : |
| http://userweb.molbiol.ox.ac.uk/public/telenius/NGseqBasicManual/external/HUBtutorial.pdf |
| UCSC genome browser data hub load instructions : |
| https://genome.ucsc.edu/goldenPath/help/hgTrackHubHelp.html#View |
| Run setup files for the above runs : |
| DNaseI+ CTCF data hub |
| http://userweb.molbiol.ox.ac.uk/public/telenius/NGseqBasicManual/external/GEODataHubs/runParameters/dnaseI_and_CTCF.tar |
| CTCF Cut-and-Run -- maxins 120 |
| http://userweb.molbiol.ox.ac.uk/public/telenius/NGseqBasicManual/external/GEODataHubs/runParameters/maxins120_GSM2433142.tar |
| CTCF Cut-and-Run -- maxins 700 |
| http://userweb.molbiol.ox.ac.uk/public/telenius/NGseqBasicManual/external/GEODataHubs/runParameters/maxins700_GSM2433142.tar |

The data hubs generated by NGseqBasic contain all the visualisations shown in the figures above, as well as provide access to all the quality control steps and original run output and error log files.

These visualisation and data metrics data hubs, and the associated NGseqBasic run setup files, can be found in Table 1, along with the instructions to load these data hubs to the USCS server.

## IMPLEMENTATION

NGseqBasic can analyse both single end and paired end fastq files, or bam files (Figure 1). Reads are mapped in either Bowtie1 ^23^ or Bowtie2 ^24^, and non-mapping reads continue to 3-prime trimming via trim_galore ^25^. The reads which don’t map after trimming, are subjected to short-read rescue via flash ^26^, which produces longer reads by combining the overlapping part of read1 and read2 of the paired end reads. All these steps are coupled with fastQC ^27^ reports, to monitor the trimming and read combining quality. For paired end sequencing, the un-mapping read pairs are further mapped as single end reads, to generate quality-control visualisations.

The properly paired reads (or all mapped reads for single end sample) continue to duplicate filtering in samtools ^28^. The data is also filtered for blacklisted regions for mm9, mm10, hg18 and hg19 genomes (additional blacklists can be provided for other genomes). The process is accompanied with samtools statistics and flagstat runs, providing simple read counts of the data before and after filtering (Figure 2).

After filtering, the sequenced fragments are reconstructed from Read1 and Read2 mapped fragments (if paired end sample) by using bedtools ^29^, and visualised in a UCSC data hub: PCR duplicates are show in red, non-duplicates in green (Figure 3). If requested, the footprints - DNaseI cut sites - are generated and plotted as well (Figure1, Figure3).

The bam file of the properly paired filtered reads is saved in the analysis folder. Other bam files (the unfiltered bam file, single end mapped files) can be saved if requested (Figure 1).

If requested, a contig-style peak call is produced (based on read depth, peak width, and distance to neighbouring peaks). The peaks are added to the visualisation data hub (Figure 3), and a multi-image-genome MIG ^30^ - loadable GFF3 file containing peak width and signal depth is generated, to flexibly assess and filter the called peak set. The fast peak call (15min run time) can be run several times, with different parameters, until satisfactory.

All the output data files are stored in the run folder, readily available for further analyses of the data set. The web link to the visualisation UCSC data hub, is given in the end of the run log file. When inspecting the data hub in UCSC browser, the analysis quality metrics and log can be readily accessed by clicking any of the tracks, to open the “track description” window (Figure 2).

The data hub can be loaded to any UCSC server instance, allowing comparisons to publicly available datasets. The hub can be accessed anywhere, simply by sharing the data hub address within the research group or with external collaborators. However, caution should be taken, as the visualisation data is not password-protected.

## AVAILABILITY, DOWNLOAD, AND HELP

Download, setup and help instructions available in the NGseqBasic web site http://userweb.molbiol.ox.ac.uk/public/telenius/NGseqBasicManual/external/

NGseqBasic can be downloaded as a tar.gz or .zip archive from https://github.com/Hughes-Genome-Group/NGseqBasic/releases. Bioconda users can load the tool as library “ngseqbasic”.

After downloading the code, follow the instructions in the tool website http://userweb.molbiol.ox.ac.uk/public/telenius/NGseqBasicManual/external/ to set up the configuration files for (i) the required toolkits (table1), and (ii) bowtie1/2 indices. You also need a (iii) public server address, to generate a data hub visible to UCSC servers. A helper script testEnvironment.sh (included in the code download) is provided to assist in the setup.

NGseqBasic is distributed under GPL3 licence.

## SUMMARY

NGseqBasic is an easy-to-use one-command tool for ATAC/DNaseI/ChIP sequencing data analysis, used in several peer-reviewed publications ^1,2,18,19^ providing 1) mapping 2) trimming 3) short read rescue 4) peak call 5) footprint 6) collecting run logs and data quality statistics and 7) one-click loadable UCSC visualisation data hub containing all the visualisation tracks, run logs and data quality statistics.

The tool provides an easy access to basic analysis, statistics, and quality metrics of a data set. All results are combined to a single UCSC data hub, enabling direct comparison to already existing public data sets. The data hub and its associated statistics html page can be accessed anywhere, simply by sharing the data hub address within the research group or with external collaborators.

NGseqBasic has been routinely used in next generation sequencing data analysis for four years, making it a robust and mature tool. It is accompanied with a version control repository, and is distributed under GPL3 licence.

## ACKNOWLEDGEMENTS

This work was supported by the Wellcome Trust Strategic Award, reference 106130/Z/14/Z) and by Medical Research Council Core Funding. We acknowledge the CBRG (Computational Biology Research Group) of Weatherall Institute of Molecular Medicine, Oxford, for maintenance and upgrading of the computational environment and cluster environment, within which NGseqBasic was developed and tested, and is still used intra-house. Especially we thank the system administrators Ewan Mac Mahon and Zong-Pei Han, as well as Simon McGowan for the core of fastqScores.pl script.

We acknowledge Agata Wesolowska-Anderson and Jason Torres (Wellcome Trust of Human Genetics, Oxford), as well as Charlotte George, Sebastian Luna Valero, and David Sims (CGAT, Computational Genomics and Analysis Training, Oxford), for testing the tool setup and installation as external users. Especially we want to thank Sebastian Luna Valero for building the BioConda support for NGseqBasic.

We acknowledge all the intra-house pipeline users for the past 4 years, and external users for the past 2 years, for bug reports and user feedback, especially Maria Suciu, and Jessica Davies, who were the long term alpha and beta testers of NGseqBasic, and Agata Wesolowska-Anderson and Jason Torres who were the alpha testers of the portable version of NGseqBasic.

---

**Table 2.** List of toolkits NGseqBasic requires.

| Toolkit | Tested with | Supports | Does not support |
| --- | --- | --- | --- |
| samtools | 0.1.19 | 0.x | 1.x |
| bowtie | 1.0.0 | 0.x , 1.x |  |
| bowtie2 | 2.1.0 | All |  |
| bedtools | 2.17.0 | 2.1x | 2.2x |
| ucsc tools | 1 | 1.x | 2.x |
| flash | 1.2.8 | 1.x | ? |
| trim_galore | 0.3.1 | 0.x | ? |
| cutadapt | 1.2.1 | 1.x | ? |
| fastqc | 0.11.4 | 0.11.x | 0.10.x |
| perl | 5.18.1 | 5.x | ? |

## REFERENCES

1. Carrat, G. R. et al. Decreased STARD10 Expression Is Associated with Defective Insulin Secretion in Humans and Mice. Am. J. Hum. Genet. 100, 238–256 (2017).

2. Hay, D. et al. Genetic dissection of the α-globin super-enhancer in vivo. Nat. Genet. 48, 1–12 (2016).

3. Guzman, C. & D’Orso, I. CIPHER: A flexible and extensive workflow platform for integrative next-generation sequencing data analysis and genomic regulatory element prediction. BMC Bioinformatics 18, 1–16 (2017).

4. H Backman, T. W. & Girke, T. systemPipeR: NGS workflow and report generation environment. BMC Bioinformatics 17, 388 (2016).

5. Fisch, K. M. et al. Omics Pipe: A community-based framework for reproducible multi-omics data analysis. Bioinformatics 31, 1724–1728 (2015).

6. Qin, Q. et al. ChiLin: a comprehensive ChIP-seq and DNase-seq quality control and analysis pipeline. BMC Bioinformatics 17, 404 (2016).

7. Yan, H. et al. HiChIP: a high-throughput pipeline for integrative analysis of ChIP-Seq data. BMC Bioinformatics 15, 280 (2014).

8. Ewels, P., Krueger, F., Kaeller, M. & Andrews, S. Cluster Flow: A user-friendly bioinformatics workflow tool [version 2; referees: 3 approved]. F1000Research 5, (2017).

9. Cormier, N. et al. Reusable, extensible, and modifiable R scripts and Kepler workflows for comprehensive single set ChIP-seq analysis. BMC Bioinformatics 17, 270 (2016).

10. Esse, R. ChIPdig: a comprehensive user-friendly tool for mining multi-sample ChIP-seq data. bioRxiv (2017). doi:https://doi.org/10.1101/220079

11. Kim, T., Seo, H. D., Hennighausen, L., Lee, D. & Kang, K. Octopus-toolkit: a workflow to automate mining of public epigenomic and transcriptomic next-generation sequencing data. Nucleic Acids Res. 46, 0–5 (2018).

12. Rubio-Camarillo, M., Gómez-López, G., Fernández, J. M., Valencia, A. & Pisano, D. G. RUbioSeq: A suite of parallelized pipelines to automate exome variation and bisulfite-seq analyses. Bioinformatics 29, 1687–1689 (2013).

13. Wagle, P., Nikolić, M. & Frommolt, P. QuickNGS elevates Next-Generation Sequencing data analysis to a new level of automation. BMC Genomics 16, 1–8 (2015).

14. Meyer, L. R. et al. The UCSC Genome Browser database: Extensions and updates 2013. Nucleic Acids Res. 41, 64–69 (2013).

15. Thurner, M. et al. Integration of human pancreatic islet genomic data refines regulatory mechanisms at type 2 diabetes susceptibility loci. Elife 7, 1–30 (2018).

16. Thurner, M. et al. Genes Associated with Pancreas Development and Function Maintain Open Chromatin in iPSCs Generated from Human Pancreatic Beta Cells. Stem Cell Reports 9, 1395–1405 (2017).

17. Hanssen, L. L. P. et al. Tissue-specific CTCF-cohesin-mediated chromatin architecture delimits enhancer interactions and function in vivo. Nat. Cell Biol. 19, 952–961 (2017).

18. Simon, C. S. et al. Functional characterisation of cis-regulatory elements governing dynamic Eomes expression in the early mouse embryo. Development 144, 1249 LP–1260 (2017).

19. Godfrey, L. et al. MLL-AF4 binds directly to a BCL-2 specific enhancer and modulates H3K27 acetylation. Exp. Hematol. 47, 64–75 (2017).

20. Neph, S. et al. An expansive human regulatory lexicon encoded in transcription factor footprints. Nature 489, 83–90 (2012).

21. Stanton, K. P., Jin, J., Lederman, R. R., Weissman, S. M. & Kluger, Y. Ritornello: high fidelity control-free chromatin immunoprecipitation peak calling. Nucleic Acids Res. 45, 1–18 (2017).

22. Skene, P. J. & Henikoff, S. An efficient targeted nuclease strategy for high-resolution mapping of DNA binding sites. Elife 6, 1–35 (2017).

23. Langmead, B., Trapnell, C., Pop, M. & Salzberg, S. L. Ultrafast and memory-efficient alignment of short DNA sequences to the human genome. Genome Biol 10, (2009).

24. Langmead, B. & Salzberg, S. L. Fast gapped-read alignment with Bowtie 2. Nat Methods 9, (2012).

25. Krueger, F. (Babraham B. Trim_galore - wrapper around cutAdapt toolkit. (2012).

26. Magoč, T. & Salzberg, S. L. FLASH: fast length adjustment of short reads to improve genome assemblies. Bioinformatics 27, 2957 (2011).

27. Andrews, S. (Babraham B. FastQC - quality control software for FASTQ files. (2010).

28. Li, H. et al. The Sequence Alignment/Map format and SAMtools. Bioinformatics 25, 2078–2079 (2009).

29. Quinlan, A. R. & Hall, I. M. BEDTools: A flexible suite of utilities for comparing genomic features. Bioinformatics 26, 841–842 (2010).

30. McGowan, S. J., Hughes, J. R., Han, Z.-P. & Taylor, S. MIG: Multi-Image Genome viewer. Bioinformatics 29, 2477 (2013).

